# Cholinergic blockade reveals a role for human hippocampal theta in memory encoding but not retrieval

**DOI:** 10.1101/2025.05.12.653487

**Authors:** Tamara Gedankien, Jennifer Kriegel, Erfan Zabeh, David McDonagh, Bradley Lega, Joshua Jacobs

## Abstract

Cholinergic dysfunction is a hallmark of Alzheimer’s disease and other memory disorders. Yet, the neurophysiological mechanisms linking cholinergic signaling to memory remain poorly understood. In this study, we administered scopolamine, a muscarinic cholinergic antagonist, to neurosurgical patients with intracranial electrodes as they performed an associative recognition memory task. When scopolamine was present at encoding, we observed disruptions to hippocampal slow theta oscillations (2–4 Hz), with selective impairments to recollection-based memory. However, when scopolamine was present during retrieval alone, we observed disruptions to slow theta without impaired memory performance. These disruptions included dose-dependent reductions in theta power, theta phase reset, and encoding–retrieval pattern reinstatement. Together, our results challenge the notion that theta oscillations are necessary for memory retrieval, and instead suggest that theta universally reflects an encoding-related neural state. These findings motivate updates to current models of acetylcholine’s role in memory and may inform future therapies targeting rhythmic biomarkers of memory dysfunction.

## Introduction

Decades of research have established a strong link between memory and the cholinergic system, which comprises the network of neurons that produce, release, and respond to acetylcholine. Evidence from neuroimaging^1–4^, post-mortem studies^5,6^, and genetic research^7^ demonstrate that Alzheimer’s disease (AD) and other memory disorders are marked by widespread loss of cholinergic pathways. Established therapies for these conditions include cholinergic agonists to mitigate memory deficits^8,9^. In experimental settings, cholinergic antagonists impair memory in both animals^10–12^ and humans^13–18^. However, the neurophysiological mechanisms through which the cholinergic system influences memory during encoding and retrieval remain largely unknown, especially in humans. This knowledge gap creates a barrier to the development of new treatments for AD and other memory disorders associated with cholinergic imbalances.

In a previous study^18^, we administered scopolamine—a cholinergic blocker targeting muscarinic acetylcholine receptors—to neurosurgical patients during a memory task, and found that memory impairment was linked to disruptions in the amplitude and timing of hippocampal theta oscillations (2–10 Hz) during encoding. Theta power in the human hippocampus has been previously linked to successful episodic memory encoding^19,20^, and our findings provided evidence that the power and phase of hippocampal theta are cholinergic-sensitive and, as such, potential therapeutic targets for memory disorders like AD.

However, in that initial study, our task design did not allow us to distinguish whether scopolamine’s effects on memory and theta activity were specific to encoding, retrieval, or both. This distinction is critical, as deficits in memory encoding and retrieval are both hallmarks of advanced AD and other cholinergic-linked disorders^21–23^. Notably, prior behavioral studies suggest that cholinergic blockade during retrieval alone is not sufficient to cause memory impairments^17,24–27^. Yet, computational models posit that cholinergic tone influences theta oscillations across both encoding and retrieval^28–30^. These models propose that acetylcholine dynamically gates hippocampal network states: promoting synaptic plasticity and theta rhythm alignment during encoding, while facilitating reactivation of stored information during retrieval through reduced cholinergic tone. Thus, cholinergic modulation was proposed as a mechanism for toggling the hippocampus between encoding and retrieval states^31–33^. To date, there has been no empirical evidence probing the neurophysiological effects of cholinergic modulation at retrieval in humans to test these models.

To address this question, we conducted an experiment combining human intracranial recordings with cholinergic modulation to isolate the effects of scopolamine at encoding versus retrieval as well as to investigate the specificity of modulation for recollection versus familiarity- and novelty-based memory. In a double-blind, within-subject design, we administered a single dose of scopolamine (or saline, in a placebo condition) to twelve neurosurgical patients as they performed an associative recognition memory task. Replicating prior behavioral findings^17,24–27^, we found no changes in memory performance when scopolamine was present during retrieval alone, but there were impairments to recollection-based memory when the drug was present prior to encoding. We then examined how scopolamine influenced three key neurophysiological markers of memory: theta power, theta phase reset, and encoding–retrieval pattern reinstatement. Despite a lack of behavioral impairments when the drug was present at retrieval alone, we found significant electrophysiological changes at retrieval, including disruptions to theta power, theta phase reset, and reinstatement of encoding-related spectral patterns. These results show that theta oscillations that appear during the recall period of a task are not essential to core memory retrieval processes, but instead could reflect other processes such as memory updating and re-encoding. We examine how these findings may promote updated theories of cholinergic tuning of the episodic memory system, with potential implications for novel therapeutic strategies.

## Results

### Cholinergic blockade during encoding, but not retrieval, impairs recollection memory

To investigate the effects of cholinergic modulation on retrieval, we administered scopolamine, a muscarinic blocker, to twelve neurosurgical patients with implanted depth electrodes while they performed an associative recognition (AR) memory task (Fig. 1a). For each subject, we recorded intracranial electroencephalographic (iEEG) signals from the hippocampus and entorhinal cortex during two randomized double-blind sessions: an experimental condition in the presence of the drug (scopolamine) and a control condition with a placebo (saline). During the encoding phase of the AR task, subjects were presented with a series of word pairs. Following a math distractor, subjects began the retrieval phase in which they were cued with another series of word pairs. For each retrieval cue word pair, subjects indicated how it related to the encoding items, in particular whether it was intact, rearranged, or new with respect to encoding (Fig. 1c). This paradigm allows us to dissociate neural correlates of recollection-based memory from recall based on familiarity and novelty detection.

**Figure 1.**
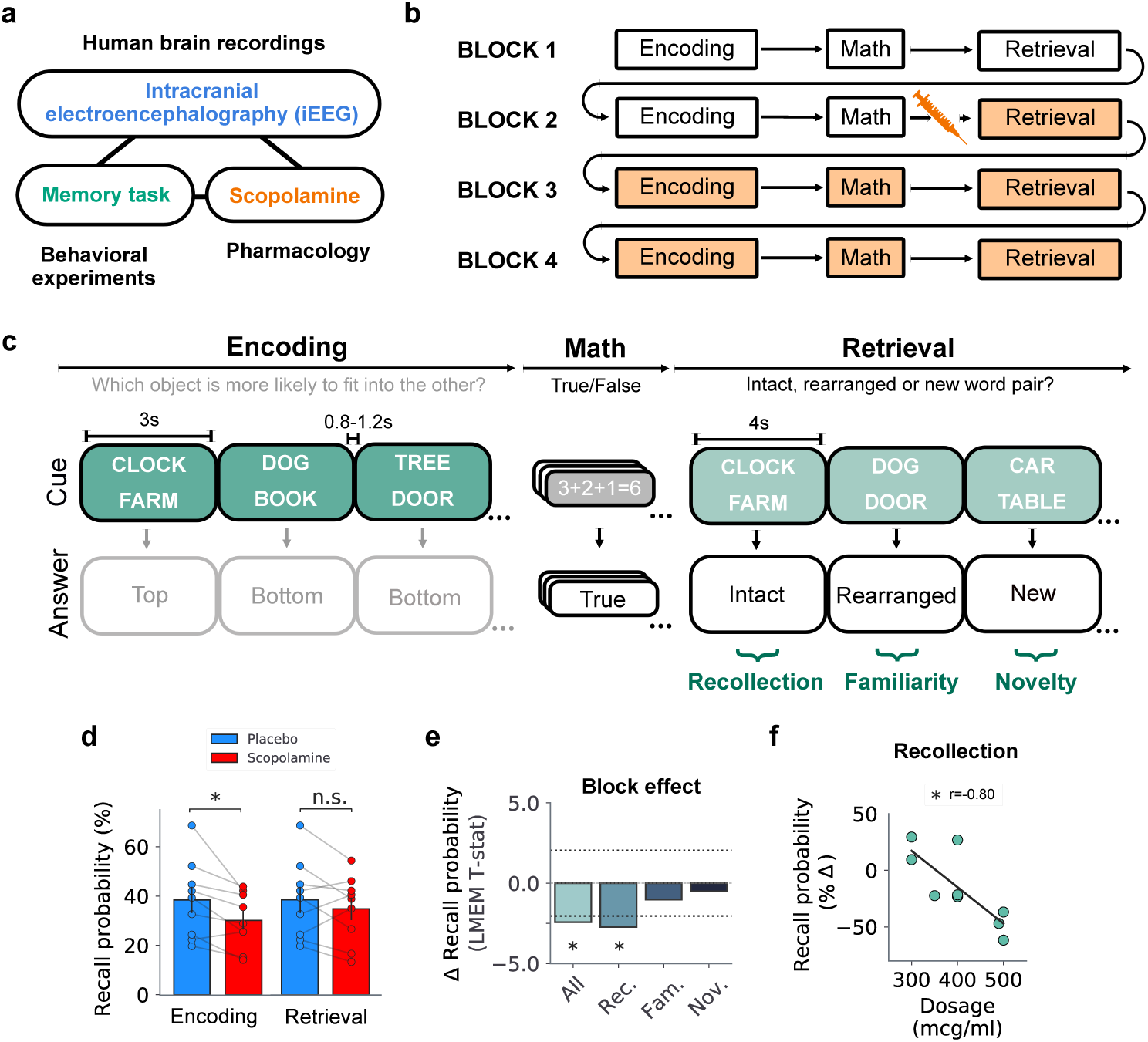
Cholinergic blockade during encoding, but not retrieval, impairs recollection memory. **a.** Overview of methods. In order to study the neural mechanisms behind the effects of cholinergic blockade on human memory, we delivered scopolamine, a cholinergic blocker, to patients with intracranial encephalographic (iEEG) electrodes during an associative recognition memory task. **b.** Diagram of the drug paradigm. In each session (drug or placebo), patients completed four blocks of the memory task. The blocks had the same task structure, starting with encoding followed by a math distractor and retrieval. In Block 2, subjects received a substance injection (scopolamine or saline), after encoding but prior to the retrieval period. Block 2’s paradigm allowed us to examine whether scopolamine impacts retrieval independently from encoding. **c.** Diagram of the task structure. Briefly, during the encoding period of the task, subjects are presented with a sequence of words pairs. Then, after completing a math distractor consisting of simple algebra equations, subjects initiate retrieval where they are asked to indicate whether word pairs presented are intact, rearranged or new in relation to the word pairs presented during encoding. **d.** Bar plot showing recall probabilities for encoding (placebo = all blocks, scopolamine = Blocks 3 and/or 4) and retrieval (placebo = all blocks, scopolamine = Block 2) across all trials. We found a significant decrease in memory performance when scopolamine is present at encoding, but not when it is present during retrieval alone (encoding Block 3 only: *t* = 42.0, *p* = 9.77*X* 10^−3^; encoding Block 4 only: *t* = 41.0, *p* = 1.37*X* 10^−2^; encoding Blocks 3 and 4: *t* = 44.0, *p* = 3.91*X* 10^−3^; retrieval: *t* = 32.0, *p* = 0.15; Wilcoxon rank sum tests). **e.** Bar plot showing change in memory as a function of block for all trials (*t*(9) = −2.42, *p* = 1.5*X* 10^−2^, LME model), and separately for recollection, familiarity, and novelty trials (recollection: *t*(9) = −2.72, *p* = 6.5*X* 10^−3^; familiarity: *t*(9) = −1.01, *p* = 0.31; novelty trials: *t*(9) = −0.50, *p* = 0.61; LME models). **f.** Correlation between percent change in memory (with respect to the placebo session) and dosage during recollection trials for Blocks 3 and 4 combined (*r* = −0.80, *p* = 0.01, two-tail Pearson’s correlation).

By using a drug paradigm with intra-task scopolamine injection, we were able to probe the effects of cholinergic blockade on encoding and retrieval separately. In each session, subjects completed four blocks of the AR task (Fig. 1b). In Block 1 (baseline), subjects completed encoding and retrieval without substance injection. In Block 2, subjects received a single injection of scopolamine (or saline, in the placebo condition) after encoding, but prior to retrieval. Thus, Block 2 allowed us to examine whether cholinergic blockade impacts retrieval independently from encoding, as individuals had encoded items in the absence of the drug but retrieved them after the injection. Lastly, subjects continued into Blocks 3 and 4, where both encoding and retrieval were affected by the substance administered in the earlier block. To assess how cholinergic blockade affected memory, we compared subjects’ accuracy between blocks and conditions (drug versus placebo), separately analyzing performance for items that required memory processes related to recollection (intact pairs), familiarity (rearranged pairs), and novelty (new pairs).

First, as expected, we found that the presence of scopolamine during encoding significantly impaired memory performance overall (placebo = all blocks, scopolamine = Blocks 3 and 4; *t* = 44.0, *p* = 3.91*X* 10^−3^; Wilcoxon rank sum test) (Fig. 1d). The effect was also significant when including Block 3 and Block 4 individually (scopolamine = Block 3 only: *t* = 42.0, *p* = 9.77*X* 10^−3^; scopolamine = Block 4 only: *t* = 41.0, *p* = 1.37*X* 10^−2^; Wilcoxon rank sum tests). However, we found no significant changes in memory when cholinergic blockade was restricted to the retrieval stage (placebo = all blocks, scopolamine = Block 2; *t* = 32.0, *p* = 0.15, Wilcoxon rank sum test) (Fig. 1d). This result is consistent with previous behavioral findings showing that cholinergic blockade during retrieval alone is not sufficient to elicit memory deficits^17,24–27^, and it was present even when excluding subjects with the lowest scopolamine doses (< 400 mcg) (Fig. S2). Notably, these effects were exclusive to items recovered on the basis of recollection rather than familiarity or novelty (recollection: *t*(9) = −2.72, *p* = 6.5*X* 10^−3^; familiarity trials: *t*(9) = −1.01, *p* = 0.31; novelty: *t*(9) = −0.50, *p* = 0.61; LME models) (Fig. 1e). Lastly, we found that changes in memory correlated with scopolamine dose during Blocks 3 and 4 (*r* = −0.80, *p* = 0.01, two-tail Pearson’s correlation) (Fig. 1f). Dosage effects were specific for items recalled on the basis of recollection, which highlights the selective impact of cholinergic blockade on memory (Fig. S1).

Overall, our results indicate that the behavioral effects of scopolamine on associative memory are selective in two ways: first, by selectively disrupting episodic memory when present during encoding but not during retrieval alone; and second, by selectively affecting hippocampal-dependent encoding processes (i.e., recollection-based memory).

### Cholinergic blockade disrupts theta power during retrieval

We next examined whether scopolamine affected electro-physiological signals at retrieval, even as memory performance remained unchanged. To test the effects of scopolamine on the amplitude of theta oscillations during retrieval, we computed changes in normalized power following the retrieval cue. For each electrode, we measured power during the entire 4-s-long retrieval period, and normalized the measured power relative to the mean and standard deviation of the baseline period 0.5 s before stimulus onset. We then compared retrieval-related shifts in slow theta power (2–4 Hz) between drug and placebo blocks.

To test whether cholinergic blockade during retrieval alone altered oscillatory activity, we analyzed data from Block 2, where subjects encoded normally but retrieved in the presence of scopolamine. We found a significant disruption in slow theta power during retrieval in the scopolamine condition compared to placebo in Block 2 (*t* = −2.93, *p* = 3.37*X* 10^−3^, LME model) (Fig. 2c). This result shows that cholinergic blockade during retrieval alone leads to theta amplitude disruption. When subjects encoded and retrieved in the setting of cholinergic blockade (Blocks 3 and 4), we also found a significant theta power disruption during retrieval in the scopolamine condition compared to placebo (*t* = −3.58, *p* = 3.46*X* 10^−4^, LME model, multiple-comparison corrected), mirroring effects observed during encoding^18^ (Fig. S4). This effect can be seen on individual electrodes, as shown in Fig. 2b, as well as at the group level, as seen in Fig. 2c. Notably, the magnitude of these effects correlated with the absolute dosage of scopolamine (*p*^′^*s <* 0.05, one-tail Pearson’s correlations) (Fig. 2d), and were present when including remembered trials only (Fig. S5).

**Figure 2.**
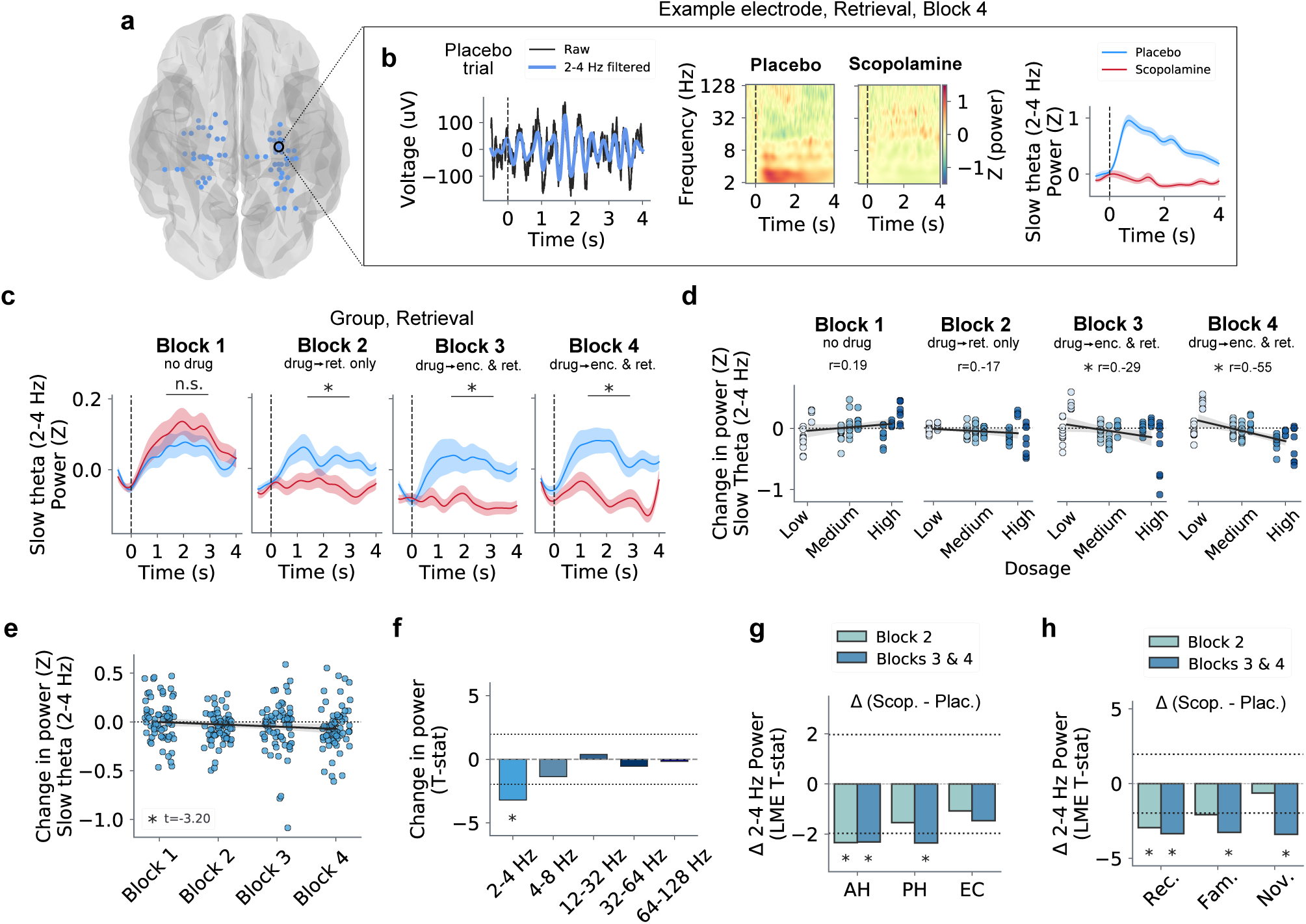
Cholinergic blockade disrupts theta power during retrieval. **a.** Anatomical locations of all hippocampal electrodes analyzed in the study. **b.** Retrieval power effects for an example electrode during Block 4. Left: line plot displaying the raw iEEG trace (black) and corresponding slow theta (2–4 Hz) filtered signal (blue) from an example trial. Middle: time-frequency spectrograms displaying mean normalized power across all trials for the example electrode during the placebo and scopolamine sessions. Time 0 s denotes the onset of the retrieval cue, and contrast denotes changes in normalized power. Right: mean slow theta power over time for placebo (blue) and scopolamine (red). Shading denotes ± SEM. **c.** Group-level mean slow theta power across all trials for Blocks 1, 2, 3 and 4 for placebo (blue) and scopolamine (red). Significant disruptions to slow theta power following scopolamine are present in Blocks 2 through 4 (*p*^′^*s <* 0.05, LME models, multiple-comparison corrected). Shading denotes ± SEM. **d.** Correlations between scopolamine dosage and change in slow theta power for all blocks (Block 1: *r* = 0.19, *p* = 0.94; Block 2: *r* = −0.17, *p* = 0.17; Block 3: *r* = −0.29, *p* = 0.02; Block 4: *r* = −0.55, *p* = 1.80*X* 10^−6^; one-tail Pearson’s correlations). **e.** Scatter plot showing change in slow theta power for all electrodes across blocks (*t* = −3.20, *p* = 1.52*X* 10^−3^, one-sided *t*-test). **f.** Change in power across blocks for different frequency bands. Only the slow theta frequency band shows a significant decrease in power across blocks. Dashed line denotes 95% significance level. **g.** Bar plot showing changes in mean slow theta power as a function of subregion (Blocks 3 and 4: anterior hippocampus (AH): *t* = −2.32, *p* = 0.02; posterior hippocampus (PH): *t* = −2.36, *p* = 0.02; and entorhinal cortex (EC): *t* = −1.46, *p* = 0.14; LME model, multiple-comparison corrected). Dashed line denotes 95% significance level. **h.** Bar plot showing changes in mean slow theta power as a function of trial category (Block 3 and 4: recollection: *t* = −3.35, *p* = 8.07*X* 10^−4^; familiarity: *t* = −3.26, *p* = 1.10*X* 10^−3^; novelty: *t* = −3.40, *p* = 6.77*X* 10^−4^; LME model, multiple-comparison corrected). Dashed line denotes 95% significance level.

Next, to better understand how power changed as a function of drug-injection paradigm, we computed the correlation between mean changes in power for each electrode across blocks. We found a significant negative correlation for changes in slow theta power across blocks (*t* = −3.20, *p* = 1.52*X* 10^−3^, one-sided *t*-test) (Fig. 2e). This correlation was only significant for the slow theta band, highlighting the particular sensitivity of the slow theta rhythm to cholinergic modulation compared to other higher-frequency oscillations (Fig. 2f).

To assess the robustness of this pattern and explore regional differences, we implemented a linear mixed-effects (LME) model that tested for differences in time-averaged theta power during retrieval. We found that power was significantly lower in the scopolamine condition compared to placebo, and this effect was driven by the anterior and posterior hippocampus (Blocks 3 and 4: anterior hippocampus (AH): *t* = −2.32, *p* = 0.02; posterior hippocampus (PH): *t* = −2.36, *p* = 0.02; and entorhinal cortex (EC): *t* = −1.46, *p* = 0.14; LME models, multiple-comparison corrected) (Fig. 2g). Lastly, we also found that changes in slow theta power during retrieval were maintained across recollection, familiarity and novelty trials (Blocks 3 and 4: all *p*’s< 0.002, LME models, multiple-comparison corrected) (Fig. 2h).

In sum, these findings indicate that cholinergic blockade changes iEEG activity during retrieval by disrupting the amplitude of slow theta oscillations, mirroring effects previously observed during encoding^18^. Importantly, despite the robust disruption to theta power during retrieval, we did not observe accompanying behavioral impairments when scopolamine was present during retrieval alone (Block 2). These results suggest the possibility that theta oscillations measured during the retrieval stage of a task may serve a function other than supporting recall processes.

### Dimensionality reduction of spectral power enables decoding of drug condition

To uncover latent structure in the data, we applied principal component analysis (PCA) to the power spectra. This unsupervised method allows us to characterize the correlational structure of retrieval-related oscillations across frequencies and brain regions, and identify features associated with pharmacological condition (Fig. 3a, b). In doing so, this data-driven analysis, which integrates data across multiple frequencies, complements our theta-focused results shown above.

**Figure 3.**
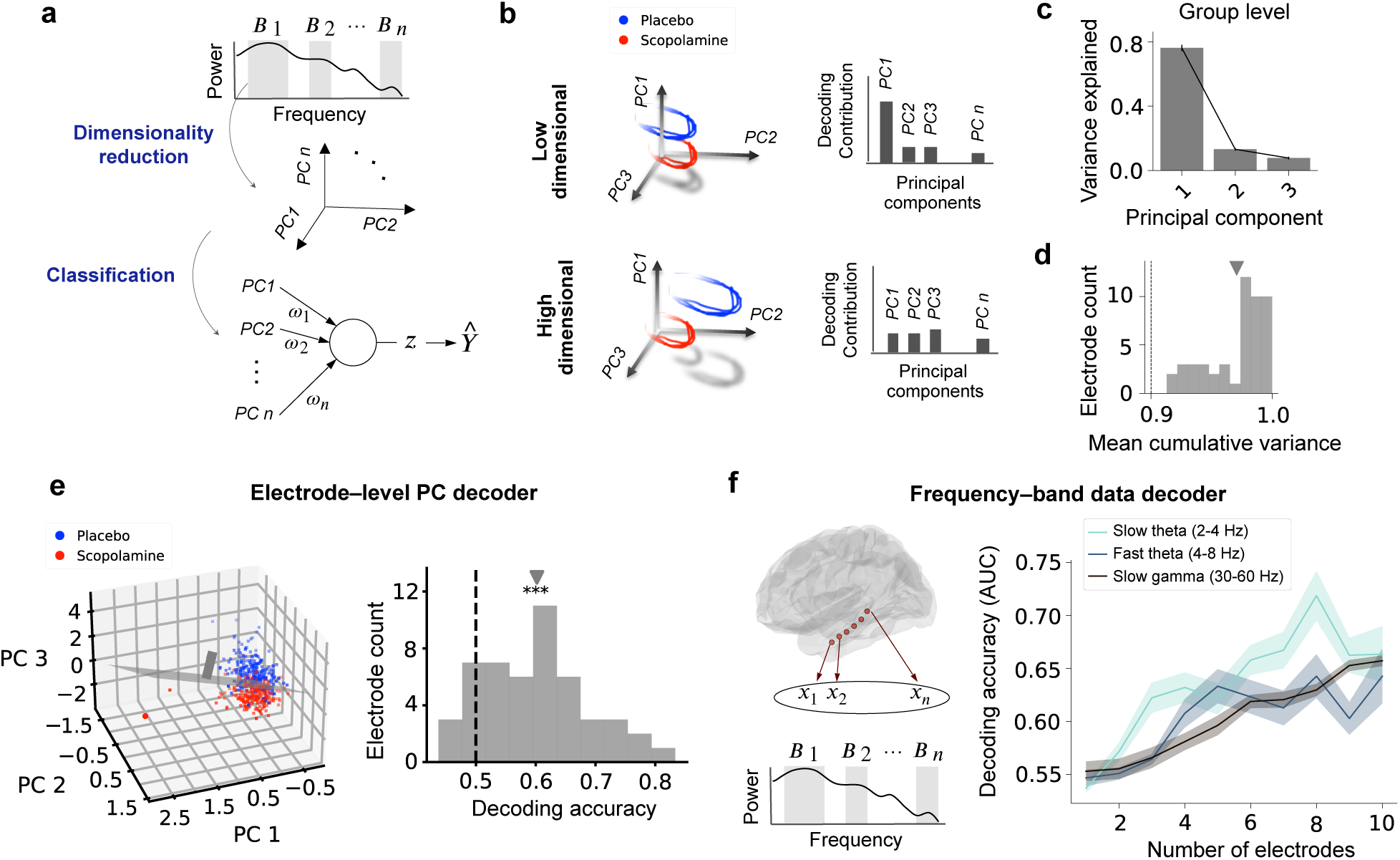
Dimensionality reduction of spectral power enables decoding of drug condition. **a.** Schematic panel outlining the analysis pipeline, including frequency-band data processing, principal component analysis (PCA) decomposition, and classification methods. This unsupervised method allows us to characterize the correlational structure of retrieval-related oscillations across frequencies and brain regions, and identify features associated with pharmacological condition. **b.** Schematic representation of low and high dimensional PCA decompositions. The separation of trajectories illustrates distinct neural representations across conditions. In low-dimensional problems, one or a few principal components (PCs) capture most of the variance and may be sufficient to support accurate decoding. In contrast, high-dimensional problems typically require many components to capture relevant variance, making decoding more complex. **c.** Bar plot showing variance explained by the first three PCs, emphasizing the dominant contribution of PC1. Error bars denotes ± SEM across electrodes. **d.** Histogram displaying mean cumulative variance (average of the first three PCs) across electrodes. The mean cumulative variance for all individual electrodes is greater than 0.9, indicating low-dimensionality in the data decomposition. **e.** Left: PCA visualization for an example electrode, showing neural trajectories from the placebo (blue) and scopolamine (red) conditions in the first three principal components. We performed dimensionality reduction for each electrode, and decoded condition (placebo or scopolamine) using the resulting PCs. Right: histogram of decoding accuracies across electrodes with a significant decoding accuracy peak at 0.6 (*p <* 0.001, Wilcoxon signed-rank test). Dashed line indicates chance-level decoding (0.5). **f.** Left: schematic of frequency-band-level decoder. We decoded condition (placebo or scopolamine) using frequency-band-filtered data across electrodes. Right: line plot comparing decoding accuracies (AUC) using slow theta, fast theta and slow gamma frequency-band-filtered data with increasing number of electrodes. Decoding accuracy increased with number of electrodes and was highest for the slow theta band, exceeding that of both fast theta and slow gamma bands.

We applied PCA to power data filtered in predefined frequency bands to identify low-dimensional electrophysiological patterns that may appear across multiple frequencies (Fig. 3a). Each resulting principal component (PC) consists of a weighted sum of power across frequency bands, capturing variability between trials. Across all electrodes, we found that the first PC accounted on average for nearly 80% of the total variance (Fig. 3c, d). The substantial variance captured by a single component suggests that, despite the complexity of the underlying neural circuits, the activity patterns relevant to memory retrieval are governed by coherent, large-scale fluctuations across frequencies and electrodes^34^.

To evaluate whether the PCA space could be used to decode pharmacological condition, we trained two linear classifiers. In the first model, we performed decoding at the electrode level, using the first three PCs as features of spectral subspace. We found that decoding accuracy was significantly above chance across all electrodes (*p <* 0.001, Wilcoxon signed-rank test) (Fig. 3e), demonstrating that the compressed PCA representation (even when using only the first PC, Fig. S8) retained sufficient information to distinguish between the placebo and scopolamine conditions. In the second model, we modified the structure and trained frequency-band-specific decoders, using electrode identity as the feature space. We found that decoding accuracy was highest for the slow theta band, exceeding that of both fast theta and slow gamma bands (Fig. 3f). Furthermore, we observed a positive relationship between decoding accuracy and the number of electrodes included in the model (Fig. 3f), suggesting that different subregions within the hippocampus contribute distinct electrophysiological signatures to the classification.

Together, these results demonstrate that scopolamine-related LFP activity can be accurately decoded from a low-dimensional feature space. Moreover, our results suggest that cholinergic blockade modulates neural dynamics along a restricted set of spectral patterns, which can enhance decoding and reflect selective engagement of dominant circuit activity, as seen other neuronal modalities^35,36^. Importantly, the strongest decoding performance emerged from the slow theta band, aligning with our hypothesis-driven findings that identified disruptions in slow theta oscillations following scopolamine administration (Fig. 2). This convergence across analytical approaches reinforces the conclusion that cholinergic modulation critically shapes slow theta dynamics during memory processing.

### Cholinergic blockade impairs theta phase alignment during retrieval

Building on evidence that the phase of ongoing neuronal rhythms facilitates memory functions^37,38^ and earlier work showing disruptions to theta phase reset during encoding following scopolamine^18^, we examined whether cholinergic blockade altered phase dynamics of theta during retrieval. We examined whether cholinergic blockade leads to changes in theta phase alignment by computing the inter-trial phase coherence (ITPC, or phase reset) at the time of the retrieval cue.

In the placebo session, we found that theta oscillations exhibited significant phase reset following the retrieval cue (Fig. 4a–c). However, the magnitude of theta phase reset was significantly reduced in Blocks 2 through 4, when the drug was present at recall (all *p*^′^*s <* 0.05, LME models, multiple-comparison corrected). This can be seen in individual electrodes as shown in Fig. 4a and b, and at the group level in Fig. 4c, where the effects were driven by the anterior and posterior hippocampus (Fig. S6). Furthermore, we found significant correlations between changes in slow theta phase reset and scopolamine dosage (*p*^′^*s <* 0.05, one-tail Pearson’s correlations) (Fig. 4d).

**Figure 4.**
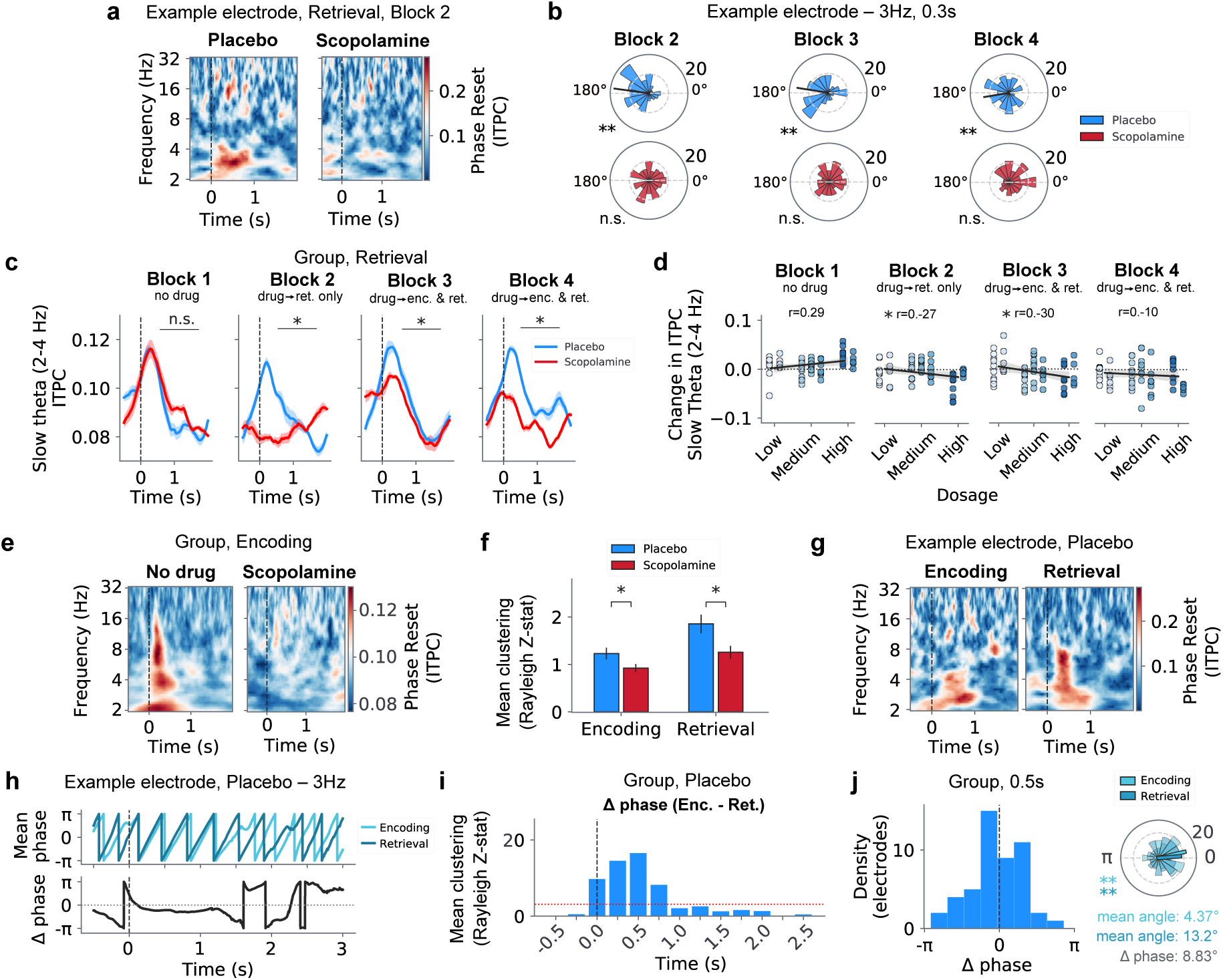
Cholinergic blockade impairs theta phase alignment during retrieval. **a.** Retrieval inter-trial phase coherence (ITPC, or phase reset) effects for an example electrode during Block 2. Time 0 s denotes onset of retrieval cue, and contrast denotes strength of phase reset. During the placebo session, the electrode shows increased slow theta (2–4 Hz) phase reset following cue onset. This effect is absent during the scopolamine session. **b.** Polar plots showing phase distribution for a 3 Hz oscillation at 0.3 s for the same example electrode. During the placebo session, the electrode exhibits significant phase clustering during Blocks 2, 3 and 4 (Block 2: *z* = 14.9, *p* = 2.27*X* 10^−7^, Block 3: *z* = 9.16, *p* = 9.15*X* 10^−5^, Block 4: *z* = 4.63, *p* = 9.54*X* 10^−3^; Rayleigh tests), but that effect is absent during the scopolamine session (Block 2: *z* = 0.23, *p* = 0.79, Block 3: *z* = 0.51, *p* = 0.60, Block 4: *z* = 2.78, *p* = 0.06; Rayleigh tests). **c.** Group-level mean slow theta phase reset is significantly disrupted following scopolamine in Blocks 2 through 4 (*p*^′^*s <* 0.05, LME models, multiple-comparison corrected) across trials. Shading denotes ± SEM. **d.** Correlations between scopolamine dosage and change in slow theta phase reset for all blocks (Block 1: *r* = 0.29, *p* = 0.98; Block 2: *r* = −0.27, *p* = 0.02; Block 3: *r* = −0.30, *p* = 0.01; Block 4: *r* = −0.10, *p* = 0.24; one-tail Pearson’s correlations). **e.** Group-level phase reset effects at encoding. **f.** Bar plot showing mean phase clustering across all electrodes during placebo and scopolamine for the encoding and retrieval periods. Scopolamine disrupts slow theta phase reset during both encoding and retrieval periods (encoding: *t* = −2.07, *p* = 0.04; retrieval: *t* = −2.69, *p* = 0.01; paired *t*-tests). **g.** Phase reset effects for an example electrode during placebo. Increased 2–4 Hz ITPC is present at both encoding and retrieval. **h.** Example electrode effects for a 3 Hz oscillation at 0.5 s. Top: mean phase. Bottom: corresponding phase difference. Phase difference between encoding and retrieval is constant and nearly zero between 0 and 1 s. **i.** Group-level consistency of slow theta phase differences between encoding and retrieval during placebo. Dashed line denotes Rayleigh test significance (95% confidence level). **j.** Group-level distribution of slow theta phase differences at 0.5 s (*z* = 8.05, *p* = 2.98*X* 10^−4^, Rayleigh test).

Since scopolamine disrupted theta phase in both the encoding and retrieval stages of the task (encoding: *t* = −2.07, *p* = 0.04; retrieval: *t* = −2.69, *p* = 0.01; paired *t*-tests) (Fig. 4e, f), we next examined the similarity in theta phase dynamics between encoding and retrieval under drug-free conditions. For each electrode during the placebo session, we computed the differences between the instantaneous phase at matched time points relative to the encoding and retrieval cues (Fig. 4g,h). We found significant matches in the phases between encoding and retrieval between 0 and 0.75 s of cue onset (Fig. 4i), with strongest clustering at 0.5 s post stimulus onset (*z* = 16.6, *p* = 1.49*X* 10^−8^, Rayleigh test) (Fig. 4j). This result indicates that theta phase alignment during encoding is recapitulated during retrieval. Additionally, between 0 and 0.75 s, the preferred theta phase at retrieval closely matched that of encoding, with a mean phase difference of 8.83^◦^ at 0.5 s (Fig. 4j, Fig. S7).

Taken together, our findings indicate that cholinergic blockade disrupts theta phase reset at retrieval in a manner that closely mirrors previously observed disruptions during encoding^18^. Rather than supporting separate phase regimes for different memory stages^28,29,39^, our results suggest that theta phase reset at retrieval reflects a reinstantiation of the encoding state, which has implications for our understanding of the role of theta in memory recall (see *Discussion*).

### Cholinergic blockade prevents encoding–retrieval pattern reinstatement

Previous studies, including noninvasive^40,41^ and intracranial EEG experiments^42,43^, have shown that stimulus-specific spectral representations emerge during memory encoding and are reactivated during retrieval—a phenomenon known as reinstatement. We therefore tested whether scopolamine disrupts spectral power reinstatement in the hippocampus to evaluate how cholinergic modulation shapes the neural conditions that support memory retrieval.

To assess whether scopolamine altered neural pattern reinstatement, we focused on recalled trials from the recollection category. We computed the correlation of spectral power (4–128 Hz) between the corresponding encoding and retrieval periods in 250 ms time bins (Fig. 5a). We chose a broadband frequency range of 4–128 Hz since reinstatement patterns have been observed at a wide range of frequency bands^44,45^. However, since we report changes in slow theta power between conditions, we excluded the slow theta frequency range (2–4 Hz) from the reinstatement correlations to avoid confounding effects. The resulting correlation matrix for an example electrode is shown in Figure 5a: the color contrast indicates the strength of the spectral power correlation between encoding and retrieval for each combination of time-bin pairs. This similarity matrix allows us to quantify whether spectral patterns from encoding are reinstated during matched, earlier, or later time points during retrieval.

**Figure 5.**
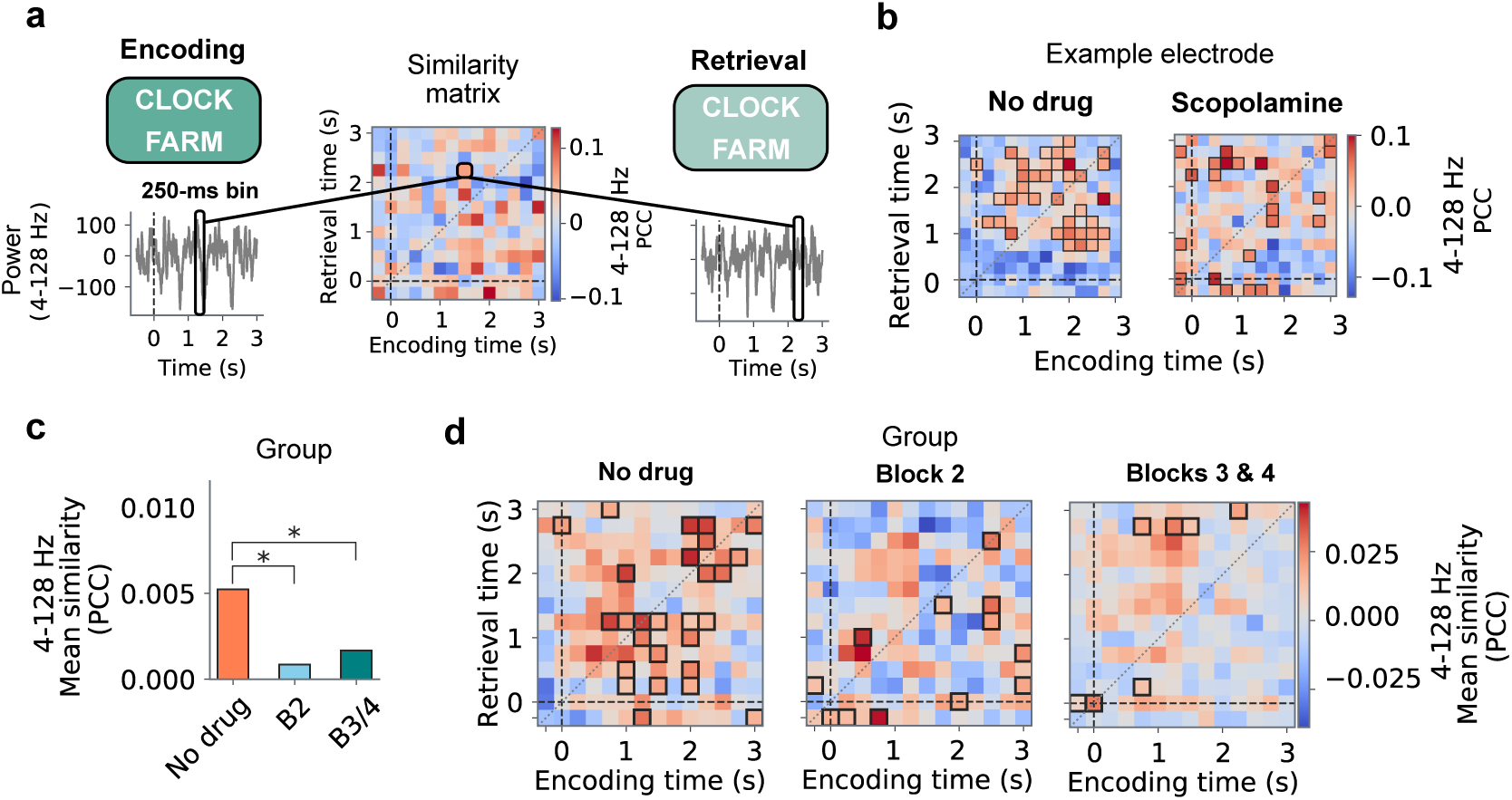
Cholinergic blockade prevents encoding–retrieval pattern reinstatement. **a.** Schematic panel outlining methods for spectral pattern reinstatement analysis. First, we identified matching encoding and retrieval periods for all recollection trials. We then computed the correlation of 4–128 Hz power between encoding and retrieval across 250 ms time bins during the initial 3 s following the encoding or retrieval cue. The resulting similarity matrix quantifies spectral reinstatement by showing whether spectral patterns at different time points during encoding are reinstated at similar, later or earlier time points during retrieval. **b.** Similarity matrix for example electrode. The “no drug” plot reflects the mean across all blocks within the placebo session, whereas the “scopolamine” plot shows combined data from Blocks 3 and 4. The contrast denotes the Pearson’s correlation coefficient (PCC) of 4–128 Hz power data between encoding and retrieval. Black outlines denote pixels exhibiting greater-than-chance reinstatement compared to surrogate data (95% confidence level). Surrogate data for electrode-level analyses involved shuffling trial labels. **c.** Bar plot showing mean reinstatement across conditions. Retrieval reinstatement during the no-drug condition is significantly greater than during the scopolamine conditions (no-drug versus scopolamine Block 2: *t* = 5.04, *p* = 1.05*X* 10^−6^; no-drug versus scopolamine Blocks 3 and 4: *t* = 5.06, *p* = 9.68*X* 10^−7^; paired *t*-tests). **d.** Group-level mean reinstatement matrices for no-drug condition, scopolamine Block 2, and scopolamine Blocks 3 and 4 combined. Black outlines denote pixels exhibiting greater-than-chance reinstatement compared to surrogate data (95% confidence level). Surrogate data for group-level analyses involved shuffling block labels.

We found that the strength of overall reinstatement was significantly greater under drug-free conditions than under scopolamine conditions. This can be seen at individual electrodes (Fig. 5b), where reinstatement is diminished during the scopolamine condition (*z* = 1.99, *p <* 0.05, two-proportion *z*-test). Similar effects appear at the group level (Fig. 5c, d), where reinstatement during no-drug conditions is significantly greater than during the scopolamine conditions (no-drug versus Block 2: *t* = 5.04, *p* = 1.05*X* 10^−6^; no-drug versus Blocks 3 and 4: *t* = 5.06, *p* = 9.68*X* 10^−7^; paired *t*-tests) (Fig. 5c). In particular, the no-drug condition shows a mean reinstatement pattern that is clustered in the matrix’s diagonal, corresponding to matched time points between encoding and retrieval. This reinstatement pattern is absent during Block 2 and Blocks 3 and 4 in the scopolamine conditions (Fig. 5d). This result indicates that cholinergic blockade disrupts both the magnitude and the timing of encoding–retrieval spectral pattern reinstatement.

Overall, these findings demonstrate that scopolamine disrupts the reinstatement of encoding-related LFP spectral patterns during retrieval, even in the absence of behavioral impairments. This suggests that while cholinergic tone may not be strictly necessary for retrieval, it may support the reactivation of an encoding-like mesoscopic neural state.

## Discussion

Although there is ample evidence that the cholinergic system is crucial for episodic memory, the specific neurophysiological mechanisms by which it modulates human memory circuits remain largely unknown. In this study, we used rare human intracranial recordings coupled with scopolamine administration to investigate how cholinergic blockade influences neuronal activity, describing neurophysiological effects during episodic memory retrieval for the first time. Replicating existing behavioral findings^17,24–27^, we showed that memory was not disrupted when scopolamine was present during retrieval alone (Fig. 1). However, when scopolamine was present during encoding, we found that it specifically impaired memory for items recovered on the basis of recollection, without impact on familiarity-based recovery or the identification of novel items. These findings are consistent with the proposed role of cholinergic circuits in mediating associative processes within the medial temporal lobe (MTL)^46,47^. We then found that scopolamine induced dose-dependent disruptions to the amplitude (Fig. 2, Fig. 3) and phase synchronization (Fig. 4) of slow theta oscillations during retrieval, including trials when the drug was present during retrieval alone, where no behavioral impairments occurred. Finally, we also showed that cholinergic blockade prevented the reinstatement of spectral patterns from encoding during retrieval (Fig. 5). Our unique results present a puzzle: why is there a disruption in an oscillatory process that supports associative memory but no commensurate change in behavior? Below, we examine how these data stimulate updated models of cholinergic influence on episodic memory processing as well as models of theta oscillations in the human MTL.

Our behavioral results extend a body of research that explored how cholinergic blockers affect behavior in both animals and humans^10–16,18^. In humans, cholinergic blockade has been shown to impair memory when present during encoding but not during retrieval alone^17,24,26,27,48^. Our results replicate these findings, as we did not find significant differences between Block 1 (the drug-free baseline) and Block 2 (when the drug was present during retrieval only). Consistent with previous work^18^, memory was significantly impaired in Blocks 3 and 4 (when the drug was present during encoding and retrieval).

Together, these results support the idea that cholinergic blockade primarily disrupts encoding but spares retrieval processes. Our behavioral results expand earlier findings by demonstrating that memory disruptions are exclusive to recollection-based memory (Fig. 2d). These findings—identified using a version of the associative recognition paradigm that is able to tease apart recollection, familiarity and novelty-based mnemonic processes—provide unique evidence for the importance of cholinergic signaling to generate item/context associations at encoding. By highlighting a selective disruption to recollection-based memory, our results shed light on potential constraints of cholinergic function in mnemonic processes. To devise treatments that address the full spectrum of memory deficiencies observed in patients with cholinergic imbalances, it will be essential to further qualify the impact of cholinergic blockade on different types of memory.

Further, our experiments directly explored how cholinergic modulation alters neurophysological processes at retrieval. Based on previous theoretical models^28,29,39^ and the lack of memory impairment following cholinergic blockade during retrieval alone^17,24–27^ (Fig. 1), we hypothesized that scopolamine would not disrupt hippocampal theta oscillatory dynamics during retrieval. On the contrary, we found that cholinergic blockade universally disrupted slow theta power (Fig. 2, Fig. 3) and phase reset (Fig. 4) during retrieval, including during trials when the drug was present during retrieval alone and encoding occurred under drug-free conditions (Block 2). Together, these observations challenge the notion that theta oscillations directly support neural processes necessary for retrieval. A key function of theta oscillations in animals^49^ and humans^50–52^ is to support the coordinated firing of neuronal ensembles on precise time scales, establishing sequences through fundamental mechanisms like spike-timing-dependent plasticity. During encoding, the importance of this process is clear: theta oscillations provide a temporal scaffold for facilitating plasticity and forming sparse neural representations, and our results are fully consistent with this model. As for retrieval, our data suggest that theta oscillations are not critical for the re-representation of those ensembles at the time of recall—because theta disruption at retrieval was not coupled with behavioral deficits. One possible synthesis of our findings with existing models is that the observed theta activity at retrieval may reflect a mechanism supporting memory updating processes that imbue flexibility and integration to representations. This potential interpretation extends earlier models (i.e., SPEAR^28,29,39^) that posit distinct theta phases for encoding and retrieval, by emphasizing that even during retrieval, the presence of theta reflects cholinergic-dependent encoding operations. This interpretation is also consistent with modern conceptions of episodic representations, which note that memory is inherently dynamic. As such, episodic memory retrieval during real-world behavior is not a passive playback of stored information, but an active, constructive process that often involves integrating the retrieval cue with ongoing contextual input.

Moreover, our PCA and decoding analyses provided new insight into how cholinergic modulation shapes memory-related neural dynamics. By showing that retrieval-related hippocampal LFP activity can be captured by a small number of principal components, we demonstrate that cholinergic disruption alters coherent, large-scale spectral patterns despite the complexity of the underlying circuits. This low-dimensional structure was sufficient to decode pharmacological condition and aligned with our hypothesis-driven results, particularly in highlighting the role of slow theta oscillations. Notably, decoding accuracy increased with the number of electrodes included, and frequency-band-specific models revealed that different electrodes contributed distinct, meaningful features to the classification. These findings suggest that theta is not a spatially uniform signal but rather exhibits regional specificity across the hippocampus, with subregions expressing unique electrophysiological signatures in response to cholinergic disruption. The superior decoding performance of the slow theta band, combined with its spatial heterogeneity, reinforces its selective vulnerability to cholinergic modulation. The convergence of unsupervised and supervised analyses reinforces the conclusion that cholinergic tone modulates hippocampal theta dynamics and highlights how local circuit activity can be leveraged to decode behavioral or pharmacological states.

Our results also include the unique finding that cholinergic blockade impairs reinstatement of oscillatory patterns at the time of retrieval. Our proposed interpretation of theta as an encoding-related signal also invites a reconsideration of the nature and functional significance of encoding–retrieval reinstatement measured at the LFP-level. Traditionally, reinstatement refers to the reactivation of neural patterns present during encoding at the time of retrieval—a mechanism thought to support successful memory recall. This property, extensively described using intracranial EEG and noninvasive imaging^40,44,45^, implies that theta synchronization, at least during encoding, supports the ability of the hippocampus to regenerate activity patterns at cue. However, if theta at retrieval reflects flexible updating of retrieved information, then recapitulation of oscillatory activity may not reflect the original memory trace in isolation, but the similarity of these theta-dependent processes between encoding and retrieval. We propose that the reinstatement signal we measure is composed of two overlapping components: one reflecting the reactivation of specific memory representations (e.g., single neurons that encoded the original experience), and another reflecting the oscillatory mechanisms that support the modification or reconsolidation of those representations. Our LFP-based reinstatement analyses lack the resolution to isolate single-neuron activity, and instead capture the broader oscillatory dynamics that support re-encoding. This distinction suggests that LFP-based reinstatement is not a direct proxy for the reinstatement of content, but rather an index of the brain’s entry into an encoding-like state that facilitates representational integration. The observed disruption to LFP reinstatement under scopolamine, then, may not reflect a failure to retrieve, but rather a failure to initiate the processes necessary for memory updating. This perspective reframes LFP reinstatement as an approximation of encoding-state reinstatement, rather than content-specific replay, and further supports our proposal that theta oscillations serve a unifying role in encoding and re-encoding across memory operations.

An alternative theory to our interpretation of theta as an encoding-related neural state, is that theta plays a role in retrieval but cued retrieval mobilizes several partly redundant representational systems, so that disruption of hippocampal theta power and phase degrades only one route to the past without fully abolishing recall capabilities. Extra-hippocampal structures such as the parahippocampal and retrosplenial cortices, posterior parietal cortex, and medial prefrontal networks have all been shown to sustain successful recall^53–55^. This redundancy is consistent with distributed frameworks in which the hippocampus orchestrates, but does not monopolize, episodic memory reconstruction^56,57^. Selective disruption of theta phase reset may therefore impair temporal sequencing while parallel extra-hippocampal pathways preserve overall retrieval performance. Even under this alternative view, our findings show that hippocampal theta is not critically required for retrieval in the same way it is for encoding.

In sum, our findings provide converging behavioral and electrophysiological evidence that cholinergic modulation plays a critical role in memory by selectively supporting hippocampal encoding processes. By showing that theta oscillations are disrupted by scopolamine during retrieval—even in the absence of behavioral impairments—we challenge the classical view that theta reflects retrieval-specific computations. Instead, we argue that theta activity that shows up at retrieval reflects an encoding-related neural mode, re-engaged during retrieval to support memory updating, contextual integration, and re-encoding. This interpretation suggest that our LFP-level encoding–retrieval reinstatement reflects the reactivation of an encoding state rather than of content alone, with implications for how we conceptualize retrieval in dynamic, reconstructive terms. More broadly, these findings prompt a reassessment of theta’s role in organizing and generating neural sequences during retrieval, and underscore the importance of distinguishing between mechanisms that support memory access and those that enable memory transformation. Future models may need to account for a role of theta oscillations in dynamically modifying previously encoded information during retrieval. Understanding how neuromodulatory systems like cholinergic pathways gate these processes will be essential for advancing both theoretical frameworks of memory and clinical strategies for memory disorders.

## Methods

### Participants

Twelve subjects participated in the study (Table S1). All participants were epilepsy patients, who received intracranial electroencephalographic (iEEG) implants as part of their treatment plan at the University of Texas Southwestern (UTSW) epilepsy surgery program.

### Ethics and safety

All participants provided their informed written consent, and the study protocol was approved by the Institutional Review Board (IRB) of Human Subjects Research of the UTSW prior to data collection. Intravenous scopolamine was administered by the participant’s attending nurse, and a board-certified anesthesiologist was present at the time of injection and remained available throughout the duration of the experiment to treat any potential adverse reactions. Physiological monitoring of blood pressure, electrocardiogram (EKG), oxygen saturation, and mental status was performed just before and for one hour after the injection of scopolamine. Low doses of scopolamine (such as 500 mcg) are commonly administered in clinical settings and generally do not present major risk factors^58^. There were no reports of adverse events related to substance administration under drug or placebo conditions.

### Scopolamine injection

Participants received a single dose of scopolamine (or saline, in the placebo session) via an intravenous line. The dosage of scopolamine for each participant was adjusted by the subject’s weight in an attempt to normalize effect sizes across subjects. The drug has a half-life of sixty-nine minutes and starts to take effect approximately five minutes after administration^58^. The second session using the alternate agent (scopolamine or saline) took place 1 to 3 days after the first session. The drug/placebo randomization was unknown to the test administrators and participants.

### Drug paradigm

Subjects completed two sessions (drug and placebo), and each session consisted of four blocks of a memory task (Fig. 1b). The blocks followed the same behavioral paradigm, but different drug paradigms. In Block 1, subjects completed the task without any substance injections. In Block 2, subjects received an injection of scopolamine or saline after encoding but prior to the retrieval period. In Blocks 3 and 4, subjects completed the task under the effects of the scopolamine (or saline) injection from Block 2 but without new interventions (Fig. 1b). This paradigm enables us to probe the effects of scopolamine on encoding and retrieval separately. In Block 2, the substance injection was administered 16 ± 2 minutes after the end of encoding, and the retrieval period commenced at 15 ± 0.2 minutes after injection (i.e., the total time between encoding and retrieval during Block 2 was approximately 31 minutes). The total session duration was 2 hours and 26 minutes ± 23 minutes, and there were no time delays between blocks beyond the substance injection procedure described above.

### Memory task

In each block, subjects completed an associative recognition (AR) memory task^59^ (Fig. 1c). During encoding, participants were shown word pairs, and later during retrieval participants were cued with word pairs that were either intact (the same as encoding), rearranged (mixed with other previously shown word pairs), or new (previously unseen words). For each retrieval cue, subjects were asked to indicate whether the cued word pair was intact, rearranged, or new. The encoding period included 90 word pairs, and each pair was displayed in capital letters for 3 seconds. To enhance and control for attention during the encoding period, after each word pair subjects were asked to indicate which of the objects in the word pair was more likely “to fit into the other”. After the encoding period, subjects completed a brief math distractor task, where they had to indicate if simple arithmetic problems in the form A + B + C = D were true or false. Next, participants began the retrieval period, which included 120 trials (60 intact, 30 rearranged, and 30 new word pairs), with cues presented for 4 seconds. During both encoding and retrieval, word pair cues were followed by an 0.8–1.2-second blank interstimulus interval (ISI) to decorrelate physiological responses from consecutive item presentations. Word pairs were randomized without repetitions across blocks.

### Intracranial recordings

Patients were implanted with either Ad-Tech Medical or PMT depth electrodes with 0.8 mm in outer diameter, forming 10 to 14 recording contacts arrayed at 4–5 mm intervals along the shaft of the electrode (Ad-Tech: 10 electrodes with variable spacing; PMT: 10 to 14 electrodes with uniform spacing, depending upon the overall depth of insertion). Intracranial EEG was sampled at 1 kHz on a Nihon–Kohden 2100 clinical system under a bipolar montage with adjacent electrodes as reference. A kurtosis threshold of 4 was applied for each trials to exclude abnormal events and interictal activity. Subject 3 was excluded from the electrophysiological analyses due to an issue with the trigger pulses. Subjects 8, 10 and 11 were excluded from the behavioral and electrophysiological analyses due to logfile errors. The electrode coverage included contacts in the posterior and anterior hippocampus, and entorhinal cortex (Table S1).

### Anatomical localization

The anatomical locations of the depth electrodes were chosen by the neurology team based on each patient’s clinical needs. Electrodes were laterally inserted into the specified regions with robotic assistance. Final electrode localization within hippocampal subregions was determined by post-operative neuroradiology review.

### Spectral power analysis

For the power analysis, we extracted spectral power using a continuous Morlet wavelet transform (wave number = 6) over 49 logarithmically spaced frequencies ranging from 2 to 128 Hz. The analysis encompassed the full 4-second retrieval periods following cue onset, with an additional 3-s buffer to minimize edge artifacts. Power values for each session, electrode, and frequency were normalized against the mean and standard deviation of the baseline period, defined as the 0.5 seconds preceding cue onset. To identify significant differences between conditions, we applied a linear mixed-effects (LME) model, using time-averaged power during the 4 s retrieval period as the dependent variable and condition (scopolamine/placebo) as a fixed effect, with electrodes treated as a random effect.

### Principal component analysis

To perform the principal component analysis (PCA), frequency-band power data for each electrode were represented as a matrix *X* ∈ R*^T^*^×*F*^, where *T* is the number of time points, and *F* is the number of spectral features. The covariance matrix of *X* was computed as:

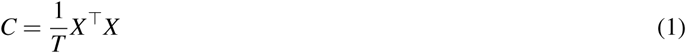

where *C* ∈ R*^F^*^×*F*^ represents the pairwise covariance between features. The eigenvectors *V* ∈ R*^F^*^×*F*^ and eigenvalues *λ* ∈ R*^F^* of *C* were then computed, where the eigenvectors define the principal components, and the eigenvalues represent the variance explained by each component. The data were projected onto the first three principal components for visualization *X*_PCA_ = *XV*_1:3_, with *V*_1:3_ containing the eigenvectors corresponding to the top three eigenvalues. The proportion of variance explained by each component was calculated as:

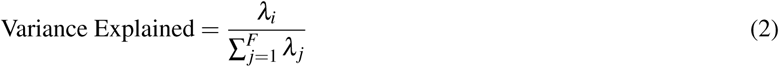

where *λ_i_* is the eigenvalue of the *i*-th principal component.

### Decoding analysis

The first decoding analysis involved an electrode-level logistic regression classifier. The input to the classifier was the power of specific frequency bands projected onto the principal components. For each trial *t*, the input features were represented as *X_t_* ∈ R*^F^*, and the binary output label *y_t_* ∈ {0, 1} indicated the drug condition. The logistic regression model was defined as:

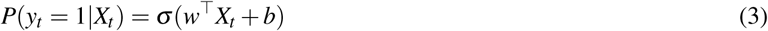

where *w* ∈ R*^F^* is the weight vector, *b* is the bias term, and *σ* (*z*) is the sigmoid function 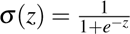. The model was trained using a cross-entropy loss function:

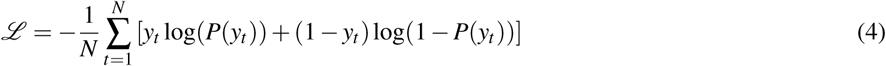

where *N* is the total number of trials. The data were divided into training (80%) and testing (20%) sets using stratified random sampling to ensure balanced class distributions. Decoding accuracies were aggregated across electrodes, and significance was determined using permutation testing with 1,000 label shuffles.

In the second decoding analysis, we performed classification using trial-wise power features from slow theta (2–4 Hz), fast theta (4–8 Hz), and slow gamma (30–60 Hz) bands to compare the discriminative power of individual frequency bands and to assess whether decoding accuracy scales with spatial coverage. For each patient, we incrementally increased the number of electrodes used in the decoder, from 1 up to the total available. At each step, we randomly sub-sampled that number of electrodes 100 times and trained a linear support vector machine (SVM) to classify trial conditions. A fixed regularization parameter (C=10) was used to prevent overfitting and dampen the influence of potentially noisy or uninformative electrodes^60^. Trial labels were balanced across conditions. All preprocessing steps, including normalization and electrode subsampling, were performed within the training data in each fold. Performance was estimated using stratified 5-fold cross-validation, and accuracy was averaged across folds and repetitions. Analyses were performed independently for each patient, and final results were averaged across patients.

### Phase reset analysis

We extracted phase measures using a continuous Morlet wavelet transformation (wave number = 6) across 49 logarithmically spaced frequencies from 2 to 32 Hz. We then computed the phase reset of oscillations at the retrieval cue onset using the inter-trial phase coherence (ITPC) measure^61,62^, which quantifies the consistency of instantaneous phase measures across trials as follows:

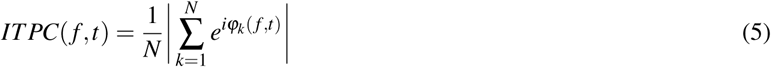

where *f* is the frequency for a given time *t*, *N* is the number of trials, and and *e^iϕk^* is the polar representation of the phase angle *ϕ*. Since ITPC calculations are susceptible to the number of observations, we excluded Subject 5, who had an incomplete number of retrieval trials (< 120) in the scopolamine condition. The ITPC was computed separately for individual electrodes, and averaged at the subject-level for the group analysis^18^.

### Reinstatement analysis

To quantify encoding–retrieval pattern reinstatement^44,63^, we first extracted 4–128 Hz spectral power in the −0.5 to 3 seconds surrounding the encoding and retrieval cues. We excluded the slow theta band (2–4 Hz), as we reported significant differences in this frequency range between conditions. Next, we binned the power data into 250-ms time bins. For each trial pair, we then computed the Pearson’s correlation of power measures between encoding and retrieval. The resulting matrix of Pearson’s correlation coefficients (PCCs) indicates the strength and direction of spectral similarity between encoding and retrieval for different time-bin combinations. Reinstatement matrices were computed separately for individual trials, and averaged at the electrode- and subject-level for the group analysis. To identify reinstatement patterns that are significantly greater than chance (95% confidence level), we compared the true distributions to surrogate data comprised of 1000 shuffle distributions. At the electrode level, surrogate data involved shuffling trial labels. At the group level, surrogate data involved shuffling block labels.

## Data availability

The data that support the findings of this study are provided in the Source Data file.

## Code availability

All data analyses were performed in Python (3.6), using publicly available software for the analysis of human electrophysiology data (https://github.com/pennmem/ptsa and https://github.com/pennmem/cmlreaders). Other custom code is available at https://github.com/tgedankien/AR_scopolamine.

## Acknowledgements

We are thankful to the patients who participated in this study. This project received support from National Institutes of Health grants R01-MH104606 (J.J.), R01-NS125250 (B.L.), and R01-NS107357 (B.L.).

## Author contributions

B.L. conceived the study and performed surgical procedures; D.M. assisted with patient screening and drug administration; J.K. and B.L. conducted the experiments; T.G., J.J., and B.L. conceived the data analyses; T.G. and E.Z. implemented the data analyses; T.G., J.J., and B.L. wrote the manuscript; J.J. and B.L jointly supervised the project.

## Supplementary Tables and Figures

**Table S1.** Subject demographics and electrode coverage. Table showing subject demographics and number of electrodes across all regions of interest, including the anterior hippocampus (AH), the posterior hippocampus (PC), and the entorhinal cortex (EC).

| Subject ID | Gender | AH | PH | EC |
| --- | --- | --- | --- | --- |
| S1 | M | 2 | 3 | 0 |
| S2 | M | 4 | 3 | 1 |
| S3 | M | 3 | 3 | 2 |
| S4 | M | 4 | 4 | 2 |
| S5 | M | 5 | 3 | 1 |
| S6 | M | 5 | 4 | 1 |
| S7 | F | 4 | 2 | 4 |
| S8 | M | 2 | 2 | 0 |
| S9 | F | 3 | 1 | 2 |
| S10 | F | 6 | 0 | 0 |
| S11 | F | 4 | 4 | 3 |
| S12 | M | 4 | 3 | 0 |

**Figure S1.**
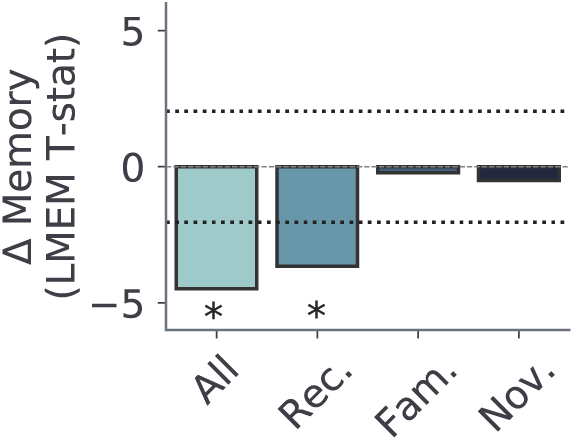
Scopolamine dosage correlates with changes in memory during recollection trials. We found significant effects of scopolamine dosage on memory. These effects were exclusive to items recovered on the basis of recollection rather than familiarity or detected novel items, consistent with a principal impact of cholinergic blockade on associative processes (all: *t* = −4.48, *p* = 7.37*X* 10^−6^; recollection: *t* = −3.65, *p* = 2.58*X* 10^−4^; familiarity: *t* = −0.22, *p* = 0.82; novelty: *t* = −0.51, *p* = 0.61; LME model, multiple-comparison corrected).

**Figure S2.**
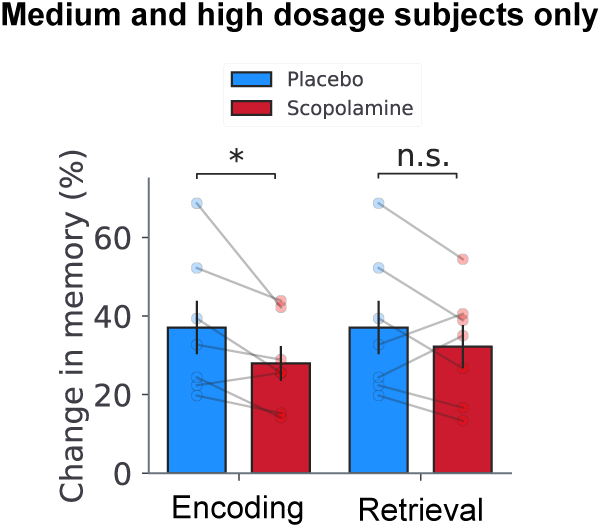
Changes in memory following scopolamine for subjects with medium and high doses only. Bar plot showing recall probabilities for encoding (placebo = all blocks, scopolamine = Blocks 3 and 4) and retrieval (placebo = all blocks, scopolamine = Block 2) across all trials for subjects with medium or high scopolamine doses (*>* 400 mcg) only. *Even after excluding subjects with lowest scopolamine doses (<* 400 *mcg)*, we found a significant decrease in memory performance when scopolamine is present at encoding, but not when it is present during retrieval alone (encoding: *t* = −2.58, *p* = 0.04; retrieval: *t* = −1.25, *p* = 0.26; paired *t*-tests).

**Figure S3.**
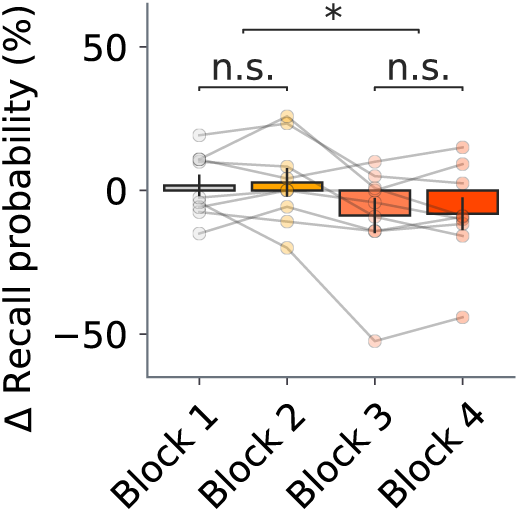
Bar plot showing change in memory across blocks for recollection trials. There are no significant differences between Block 1 (baseline) and Block 2 (when the drug is present during retrieval only) (*t* = 0.34, *p* = 0.74; paired *t*-tests). This shows that cholinergic blockade during retrieval alone is not sufficient to elicit memory deficits. However, memory is modestly worse in Blocks 3 and 4 (when the drug is present during both encoding and retrieval) compared to Blocks 1 and 2 (*t* = 2.232, *p* = 0.056, paired *t*-tests).

**Figure S4.**
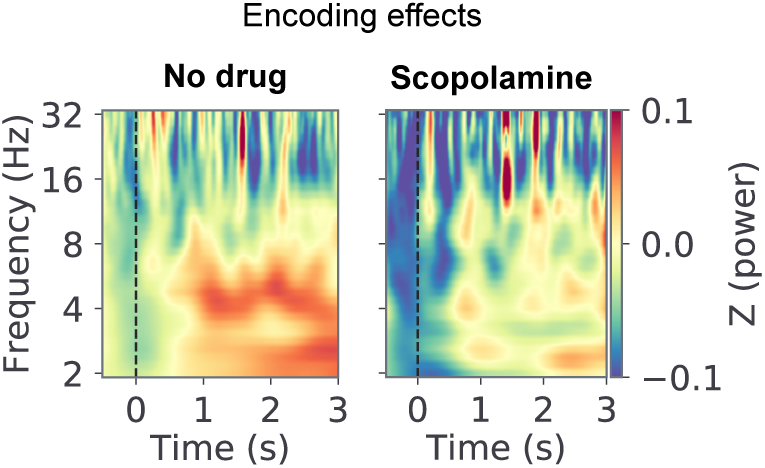
Cholinergic blockade disrupts slow theta power during encoding. Group-level time-frequency spectrograms displaying changes in mean normalized during encoding for the no-drug (all drug-free blocks combined) and scopolamine conditions. Time 0 s denotes onset of encoding cue. Contrast denotes change in normalized power. Mirroring earlier reports^18^, in the placebo session there was higher slow theta power during memory encoding, but this effect is absent during the scopolamine session.

**Figure S5.**
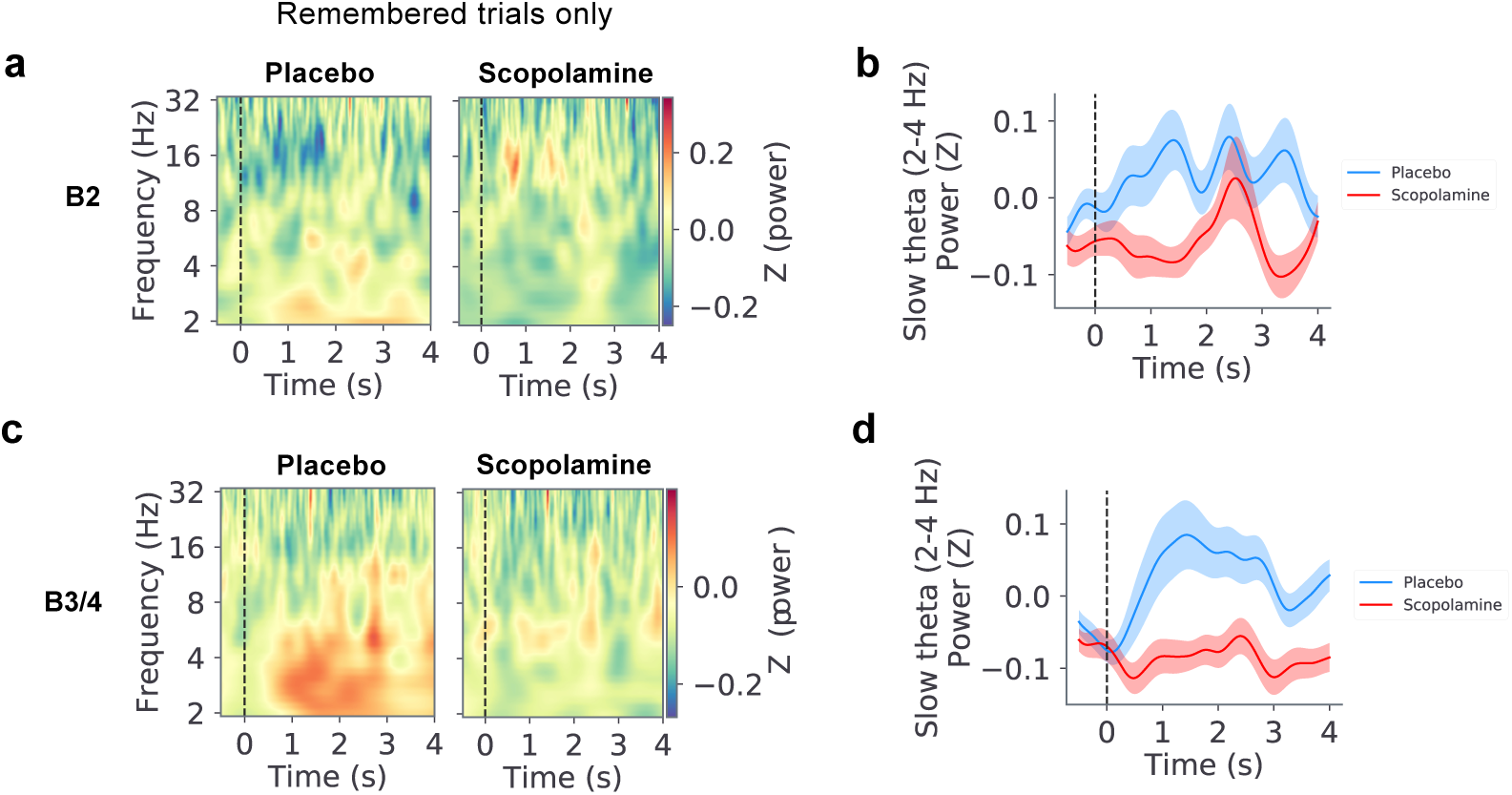
Disruptions to retrieval slow theta power in remembered trials only. Patterns of retrieval slow theta power disruptions during remembered trials are consistent with the disruptions observed across all trials (remembered and forgotten). **a.** Group-level time-frequency spectrograms displaying mean normalized for the placebo and scopolamine session during Block 2 for remembered trials only. Time 0 s denotes onset of encoding cue. Contrast denotes change in normalized power. **b.** Mean slow theta power over time for placebo (blue) and scopolamine (red) sessions for Block 2 (*p <* 0.05, LME model, multiple-comparison corrected). Shading denotes ± SEM. **c.** Group-level time-frequency spectrograms displaying mean normalized for the placebo and scopolamine session during Blocks 3 and 4 combined for remembered trials only. Time 0 s denotes onset of encoding cue. Contrast denotes change in normalized power. **d.** Mean slow theta power over time for placebo (blue) and scopolamine (red) sessions for Blocks 3 and 4 combined (*p <* 0.05, LME model, multiple-comparison corrected). Shading denotes ± SEM.

**Figure S6.**
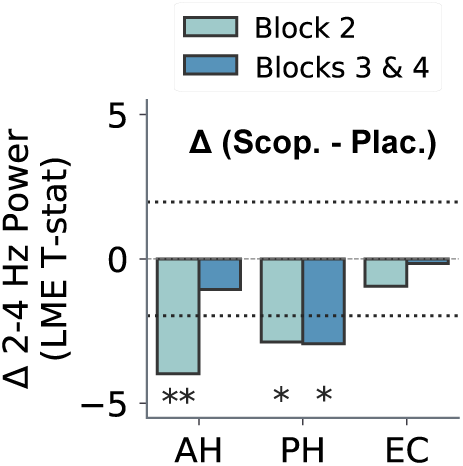
Disruptions in retrieval phase reset following cholinergic blockade are driven by the anterior and posterior hippocampus. Bar plot depicting changes in slow theta (2–4 Hz) power for different medial temporal lobe (MTL) subregions. The effects are strongest in the anterior and posterior hippocampus (Blocks 3 and 4: anterior hippocampus (AH): *t* = −3.97, *p* = 7.1*X* 10^−5^; posterior hippocampus (PH): *t* = −2.87, *p* = 4.06*X* 10^−3^; and entorhinal cortex (EC): *t* = −0.94, *p* = 0.34; LME model, multiple-comparison corrected).

**Figure S7.**
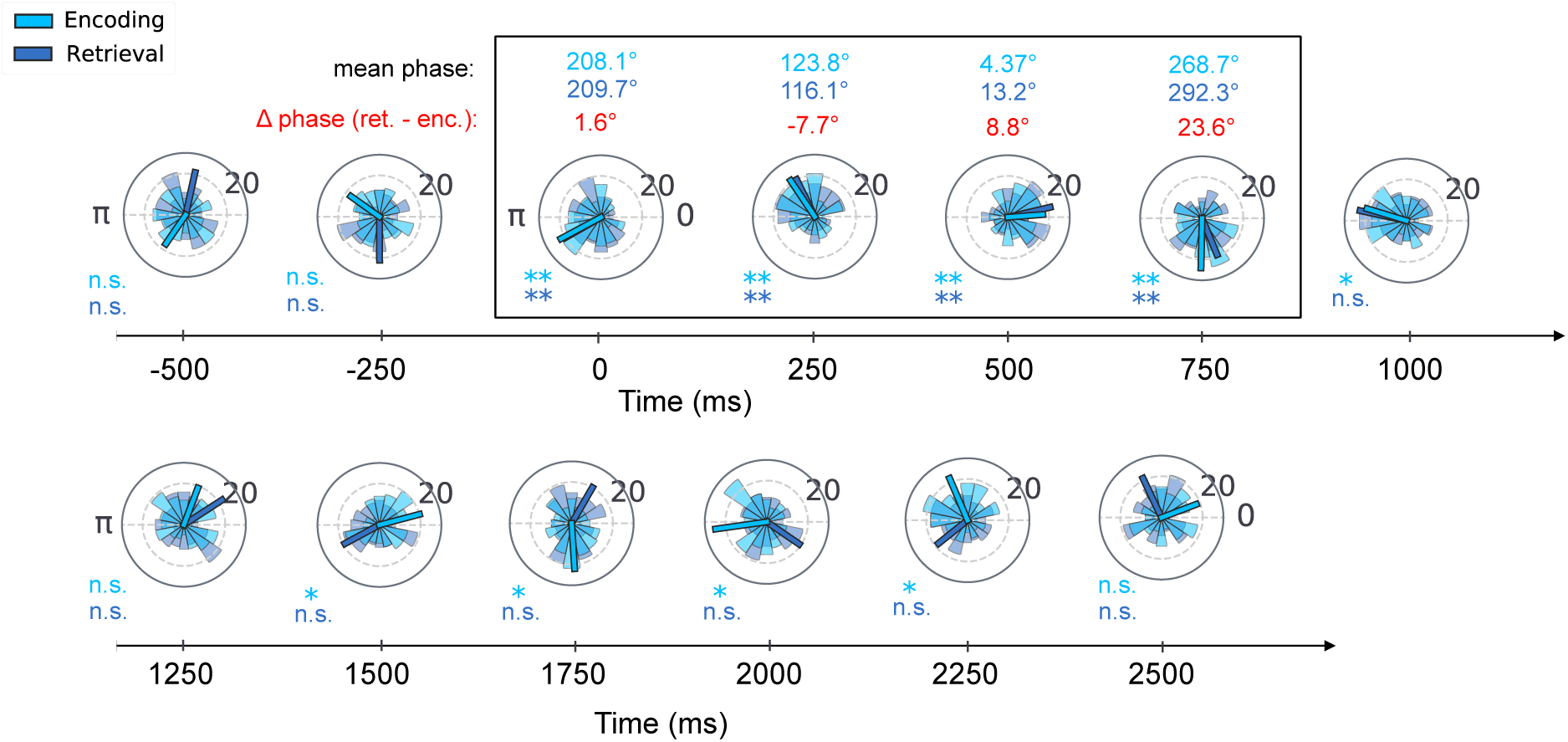
Mean preferred phase at encoding is recapitulated at retrieval. Polar plots depicting phase distributions and mean phases across all electrodes for encoding (light blue) and retrieval (dark blue). At time 0 s, the preferred phase for encoding and retrieval closely match, with mean a phase difference of only 1.6^◦^. The mean angles for encoding and retrieval remain significantly clustered until 0.75 s following the cue.

**Figure S8.**
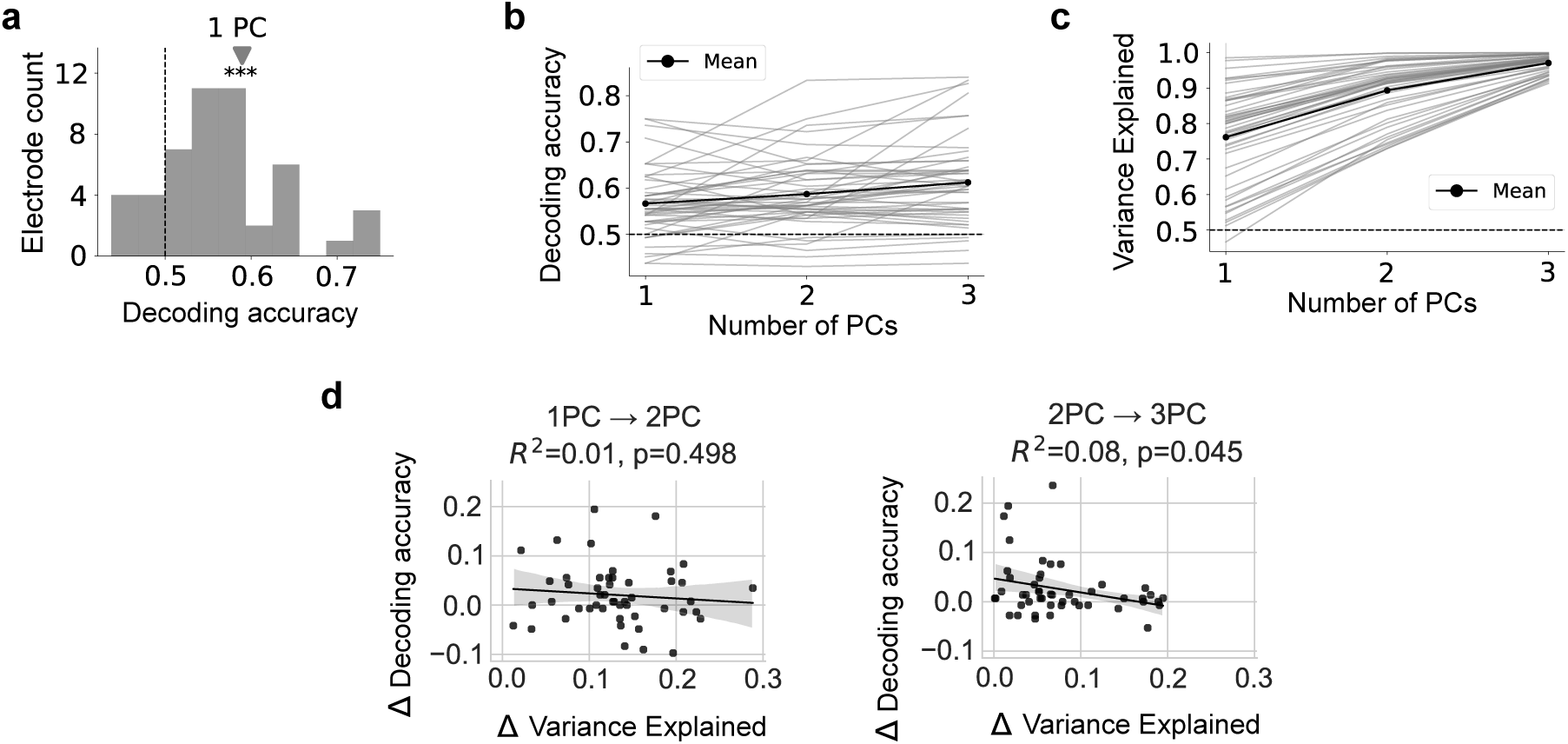
Dimensionality of spectral decoding subspace. **a.** Histogram of decoding accuracy across electrodes using only the first principal component (PC1). Mean accuracy (triangle) is significantly above chance (*p <* 0.001, Wilcoxon signed-rank test). **b.** Electrode-level decoding accuracy across PCA dimensions (1–3 PCs). While individual decoding performance increases slightly, the improvement of decoding with higher dimensionality is modest. **c.** Cumulative variance explained across PCA dimensions. Increasing the number of PCs substantially improves the variance explained, indicating that higher components capture more spectral variability but not necessarily more task-relevant information. **d.** Correlations between change in decoding accuracy and change in variance explained between 1PC and 2PCs (left), and 2PCs and 3PCs (right). The weak or absent correlation between variance gain and decoding improvement underscores that not all variance dimensions are behaviorally relevant. While additional principal components significantly increase explained variance, their contributions to decoding accuracy remain minimal or even negative. These findings support a model where early PCs capture most task-relevant structure, and later PCs primarily reflect noise or irrelevant variability.

